# scCancerExplorer: a comprehensive database for interactively exploring single-cell multi-omics data of human pan-cancer

**DOI:** 10.1101/2024.06.24.600356

**Authors:** Changzhi Huang, Zekai Liu, Yunlei Guo, Wanchu Wang, Zhen Yuan, Yusheng Guan, Deng Pan, Zhibin Hu, Linhua Sun, Zan Fu, Shuhui Bian

**Affiliations:** State Key Laboratory of Reproductive Medicine and Offspring Health, Nanjing Medical University, Nanjing 211166, China; Department of General Surgery, the First Affiliated Hospital of Nanjing Medical University, Nanjing 210029, China; College of Life Sciences, Nanjing Normal University, Nanjing 210023, China; Collaborative Innovation Center for Cancer Personalized Medicine, School of Public Health, Nanjing Medical University, Nanjing 211166, China

**Keywords:** Database, human pan-cancer, single-cell multi-omics

## Abstract

Genomic, epigenomic, and transcriptomic alterations are hallmarks of cancer cells, and are closely connected. Especially, epigenetic regulation plays a critical role in tumorigenesis and progression. The growing single-cell epigenome data in cancer research provide new opportunities for data mining from a more comprehensive perspective. However, there is still a lack of databases designed for interactively exploring the single-cell multi-omics data of human pan-cancer, especially for the single-cell epigenome data. To fill in the gap, we developed scCancerExplorer, a comprehensive and user-friendly database to facilitate the exploration of the single-cell genome, epigenome (chromatin accessibility and DNA methylation), and transcriptome data of 50 cancer types. Five major modules were provided to explore those data interactively, including “integrated multi-omics analysis”, “single-cell transcriptome”, “single-cell epigenome”, “single-cell genome”, and “TCGA analysis”. By simple clicking, users can easily investigate gene expression features, chromatin accessibility patterns, transcription factor activities, DNA methylation states, copy number variations, and TCGA survival analysis results. Taken together, scCancerExplorer is distinguished from previous databases with rich and interactive functions for exploring the single-cell multi-omics data of human pan-cancer. It bridges the gap between single-cell multi-omics data and the end-users, and will facilitate progress in the field of cancer research. scCancerExplorer is freely accessible via https://bianlab.cn/scCancerExplorer.

## INTRODUCTION

Human cancers are characterized by molecular alterations in the genome, epigenome, and transcriptome with drastic heterogeneity, which hinder the effective clinical diagnosis and treatment (1–3). In addition to cancer genome and transcriptome which have been extensively studied, aberrant epigenome plays a critical role in tumorigenesis and progression by affecting gene expression and genome integrity. Moreover, epigenetic modifications are potential therapeutic targets and have raised hope for developing effective cancer therapy. Several epigenetic agents have been proven for clinical treatment or in development, such as the DNA methylation inhibitors azacitidine and decitabine (4). A better understanding of the epigenetic regulation mechanism of cancer will greatly promote the development of clinical therapeutic strategies.

Single-cell multi-omics techniques have revolutionized cancer research over recent years, offering powerful tools to dissect the intricate molecular basis of human cancers. Large amounts of single-cell multi-omics data, encompassing the genome, transcriptome, and epigenome, have been continuously generated for human cancer studies (5, 6). Especially, with advanced single-cell multi-omics sequencing techniques, we and others can even simultaneously analyze the genome, epigenome, and transcriptome of the same single cell, and reveal the complicated relationships between multiple molecular layers (7–11). These studies not only provided a comprehensive atlas of human cancers but also offered a wealth of data resources for further exploration. Notably, the growing single-cell chromatin accessibility and DNA methylation data provide new opportunities for understanding human cancer from the epigenetic perspective (7, 9–17). However, analyzing and integrating the massive single-cell multi-omics data from separate studies posed significant challenges, particularly for wet-lab biologists and oncologists lacking analytical skills and adequate computing resources. Hence, integrated and user-friendly platforms are important to bridge the gap between single-cell multi-omics data and users.

Although several databases have been developed to facilitate cancer research, there is still a lack of online databases that can meet the need to explore single-cell multi-omics data of human pan-cancer, especially for the single-cell epigenome data (such as chromatin accessibility and DNA methylation). For example, GEPIA2 is a great tool to explore and visualize the gene expression profiles of TCGA data (18), which is widely used in cancer studies, but it does not contain single-cell data. Other valuable single-cell cancer databases, such as CancerSEA (19), TISCH2 (20), SCAR (21), and CancerSCEM (22), focus on the single-cell transcriptome and spatial transcriptome of cancer, and lack single-cell epigenome and multi-omics integration. Although we and other groups have published several single-cell epigenome or multi-omics databases, such as SMARTdb (23), SCA (24), scMethBank (25), and scATAC-Ref (26), none of them focus on human cancers. In summary, the practical needs of cancer researchers, such as exploring single-cell ATAC-seq and DNA methylation data, and acquiring integrated multi-omics information, cannot be met by the existing databases.

Here, we introduce scCancerExplorer (https://bianlab.cn/scCancerExplorer), an integrated database designed to provide user-friendly functions for exploring single-cell multi-omics data of human pan-cancer. By facilitating data mining and inspiring novel insights into pan-cancer multi-omics data, scCancerExplorer aims to empower researchers to decipher the complexities of human cancer and pave the way for cancer research.

## MATERIALS AND METHODS

### Data collection

In total, 161 single-cell multi-omics datasets covering over 6.2 million single cells (after quality control) and 50 human cancer types have been collected and analyzed, involving single-cell transcriptome, epigenome (DNA methylation and chromatin accessibility), and genome data (copy number variation) data (Table S1). Related metadata, such as pathological subtype, sex, sample location, and clinical stage, were collected if available. The TCGA data from 33 cancer types, including the gene expression matrix, DNA methylation data generated using Illumina HumanMethylation450 BeadChip, mutation information, and clinical data were obtained from UCSC Xena (https://xenabrowser.net/datapages/?hub=https://gdc.xenahubs.net:443). The ATAC-seq data from TCGA project was obtained from https://gdc.cancer.gov/about-data/publications/ATACseq-AWG (27).

### Data processing

#### Single-cell transcriptome data processing

The single-cell gene expression matrices were processed utilizing Seurat (v.4.3.0.1) (28). Briefly, quality control of the single-cell data was firstly performed according to the description in the original study. Additionally, we have established a uniform minimum standard: gene number >500 and UMI count >1000. For all of the scRNA-seq datasets generated by 10x Genomics platform, the UMI count data were all consistently normalized using the “NormalizeData” function in Seurat with “scale.factor=10000”. The top highly variable genes (default or as the original study described) were selected using the “FindVariableFeatures” function. The normalized data were then centered and scaled using the “ScaleData” function. Principal component analysis (PCA) was conducted on the scaled data using the highly variable genes. Subsequently, the uniform manifold approximation and projection (UMAP) was performed and employed to visualize the cell clustering result. The cell type identities were annotated based on available metadata or canonical marker genes. In principle, the cell type identity information provided by the original studies and TISCH2 database (20) was preferentially used. Significant differentially expressed genes (DEGs) between various cell types were identified using the “FindAllMarkers” function in Seurat.

#### Single-cell DNA methylation data processing

For each cell, only the CpG sites (for scBS-seq data) or WCG (W = A, T) sites (for scCOOL-seq data) with DNA methylation levels < 0.1 or > 0.9 were retained. The gene promoter regions were defined as 1 kb upstream and 0.5 kb downstream of the transcription start site. For each cell, only the promoter regions with at least 3 CpG or WCG sites covered were retained, and the mean methylation levels within the promoter regions were calculated to represent the methylation status of promoter regions. To plot the DNA methylation status of each cell group, the single-cell DNA methylation data were merged by cell group. For the CpG or WCG sites covered by multiple cells of the group, the mean DNA methylation levels of covered cells were calculated.

#### Single-cell ATAC-seq data processing

To maintain maximum consistency with the original study results, only the cells present in the metadata information provided by the original study were retained. For datasets lacking metadata information, low-quality cells were removed mainly based on the criteria described in the original study. Downstream data processing was conducted using the R package ArchR (v.1.0.2) (29). For datasets without provided quality control criteria in the original study, only cells with TSS enrichment score ≥ 4 and mapped fragments ≥ 1000 were used for further analyses. Doublet removal was performed using the “addDoubletScores” and “filterDoublets” function of ArchR with default parameters. The iterative latent semantic index (LSI) dimension reduction was performed using the “addIterativeLSI” function in ArchR. Peak calling was performed on single-cell ATAC-seq data using MACS2 (v.2.2.9.1). The enrichment of TF motifs for peaks in each cell cluster was performed using chromVAR (v.1.20.2). The “getGroupBW” function from ArchR package was utilized to export bigwig files with a tile size of 100 bp based on cell type. Peak-to-peak co-accessibility information was exported using the “getCoAccessibility” function and then converted to the inter.bb format using “bedTOBigBed” (30).

#### Copy number variation data processing

The single-cell copy number variation data was provided by the authors from the original studies.

#### TCGA survival analysis

The “lifelines” library (v.0.27.8) in Python was utilized for the survival analysis. The expression levels were categorized into “High” and “Low” groups based on the median or quantile expression value of the selected gene. The Kaplan-Meier survival curves were generated using the “KaplanMeierFitter” class in Python, and the log-rank tests were performed. Additionally, a Cox proportional hazards model was fitted using the “CoxPHFitter” class in Python to estimate the hazard ratios. The survival curves and statistical results were visualized with “matplotlib” (v.3.7.5) and “seaborn” (v.0.13.2).

#### TCGA DNA methylation data processing

For TCGA HumanMethylation450 data, the annotation information regarding the methylation probes was downloaded from the Illumina (https://support.illumina.com/downloads/infinium_humanmethylation450_product_fil es.html). To calculate the promoter DNA methylation levels, the gene promoter regions were defined as spanning from 1 kb upstream and 0.5 kb downstream of the transcription start site. For each sample, the DNA methylation levels for the promoter regions are calculated using the average beta values of methylation probes that fall within the gene promoter regions.

#### Implementation of the database

The scCancerExplorer platform is developed with Python and the Django framework (https://djangoproject.com), utilizing PostgreSQL (https://www.postgresql.org) for systematic data organization and management. The user-friendly interface is built with Angular and NG-ZORRO. The platform is hosted on an Nginx web server (http://nginx.org) and is freely accessible without any need for registration or login. For the best user experience, scCancerExplorer is optimized for web browsers like Google Chrome, Microsoft Edge, Safari, and Mozilla Firefox, with JavaScript enabled.

## DATABASE CONTENTS AND USAGE

### Overall design

scCancerExplorer was designed to enable convenient exploration of single-cell multi-omics data of human pan-cancer (Figure 1) and inspire new insights into the complex landscape of human cancers. In total, multi-omics data from 161 datasets covering over 6.2 million single cells (after quality control) were curated and analyzed. We provided five major modules for users, including “Integrated multi-omics analysis”, “Single-cell transcriptome”, “Single-cell epigenome”, “Single-cell genome”, and “TCGA analysis”. Twelve useful functions were provided, which will be described in detail below.

**Figure 1.**
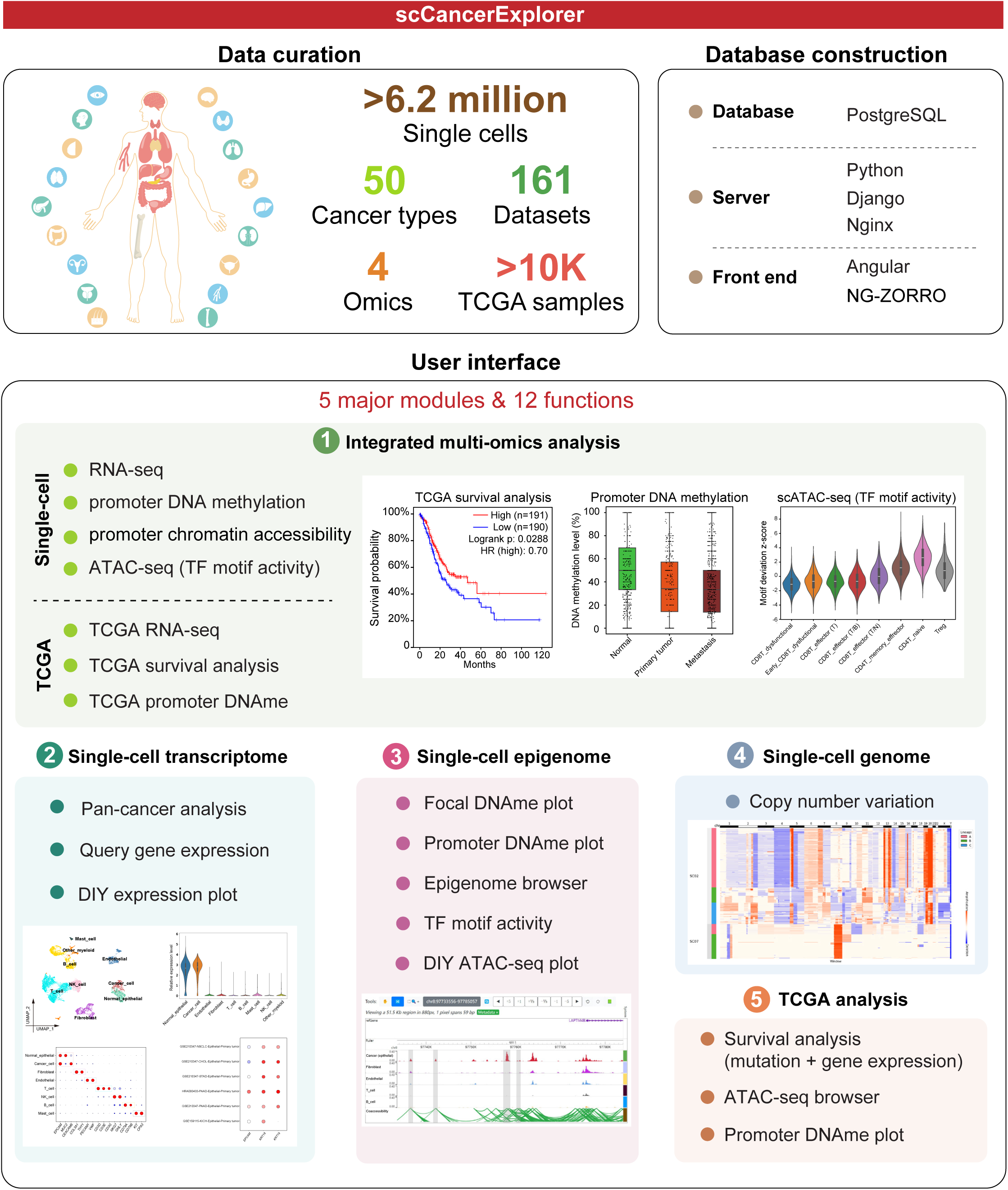
A schematic diagram illustrating the data curation, web construction, and user interface of scCancerExplorer. DNAme, DNA methylation.

### “Integrated multi-omics analysis” module

This module provided both single-cell multi-omics and TCGA multi-omics perspectives to learn an interested gene quickly (Figure 2). For example, users can explore the single-cell gene expression levels (Figure 2A), and single-cell chromatin accessibility levels and DNA methylation levels of the promoter region of the selected gene (*CMTM3*) in both normal cells and stomach cancer cells (Figure 2B, C) (9). Additionally, they can search the gene expression levels and promoter DNA methylation levels of both normal and cancer tissues using TCGA data (Figure 2D, E), as well as TCGA survival analysis results based on the expression level of the selected gene (Figure 2F). If the gene being searched is a transcription factor (TF), the TF motif activity deduced from single-cell ATAC-seq data is also provided. If users want to further learn the interested gene in depth, they can use other modules that we provided for each omics below.

**Figure 2.**
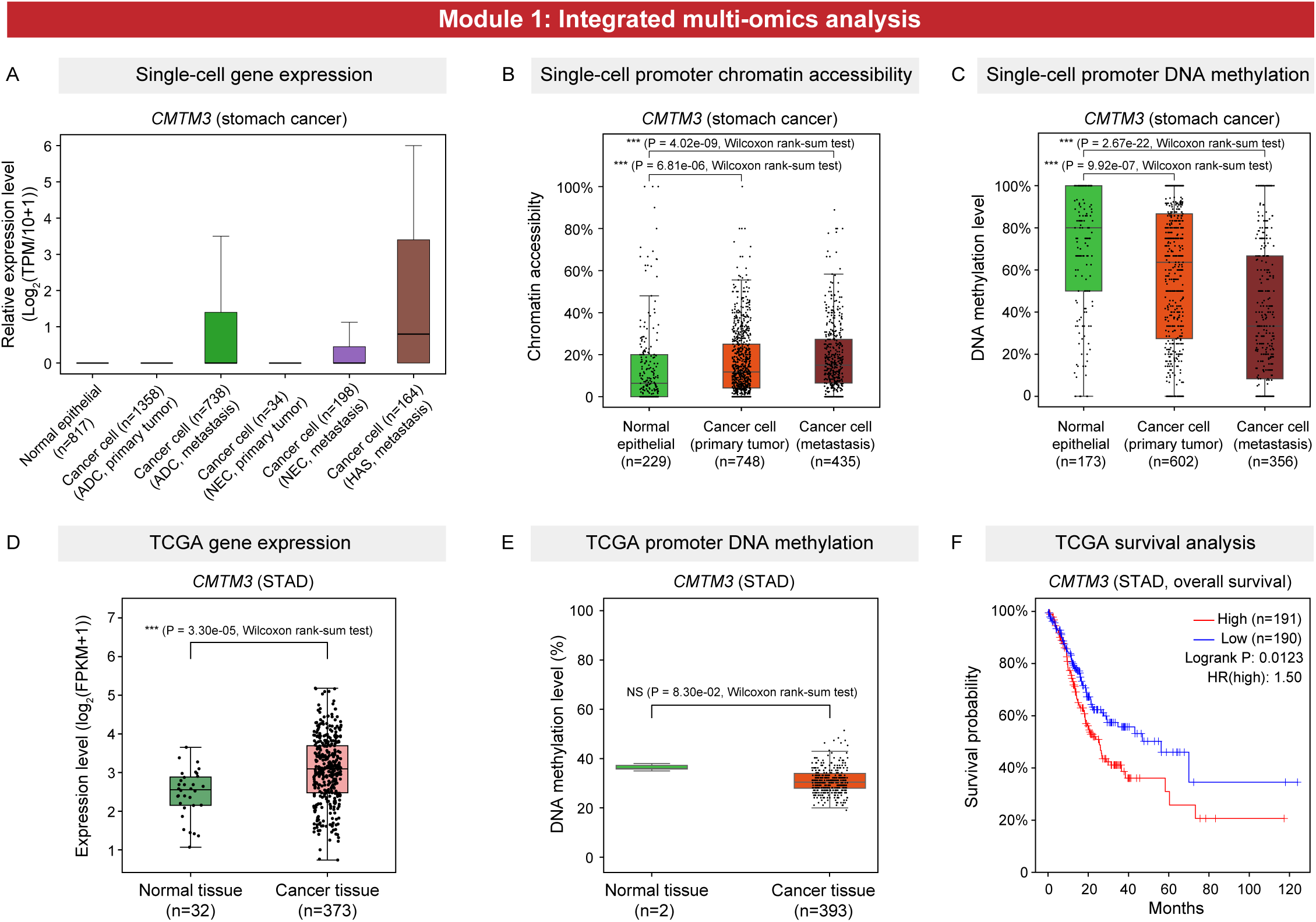
An example for exploring an interested gene with the “Integrated multi-omics analysis” module. (A-C) The single-cell gene expression levels (A), promoter chromatin accessibility levels (B), and promoter DNA methylation levels (C) of the selected gene across various cell types in stomach cancer (dataset ID: HRA001689). (D-E) The gene expression levels (D) and promoter DNA methylation levels (E) of the selected gene in normal tissues and cancer tissues in TCGA stomach adenocarcinoma dataset. (F) Overall survival analysis using TCGA stomach adenocarcinoma dataset shows that the patients with higher expression levels of the selected gene (>median value) have significantly poorer prognosis. ADC, adenocarcinoma. NEC, neuroendocrine carcinoma. HAS, hepatoid adenocarcinoma of stomach. *CMTM3*, CKLF Like MARVEL Transmembrane Domain Containing 3. STAD, stomach adenocarcinoma. NS, non-significant. ***, P value < 0.001, Wilcoxon rank-sum test.

### “Single-cell transcriptome” module

#### Pan-cancer analysis

With this function, a quick exploration of the integrated single-cell RNA-seq data of human pan-cancer is allowed (Figure 3A). For example, users can query the gene expression levels of multiple interested genes in the lymphatic endothelial cells of human pan-cancer (Figure 3A) (31).

**Figure 3.**
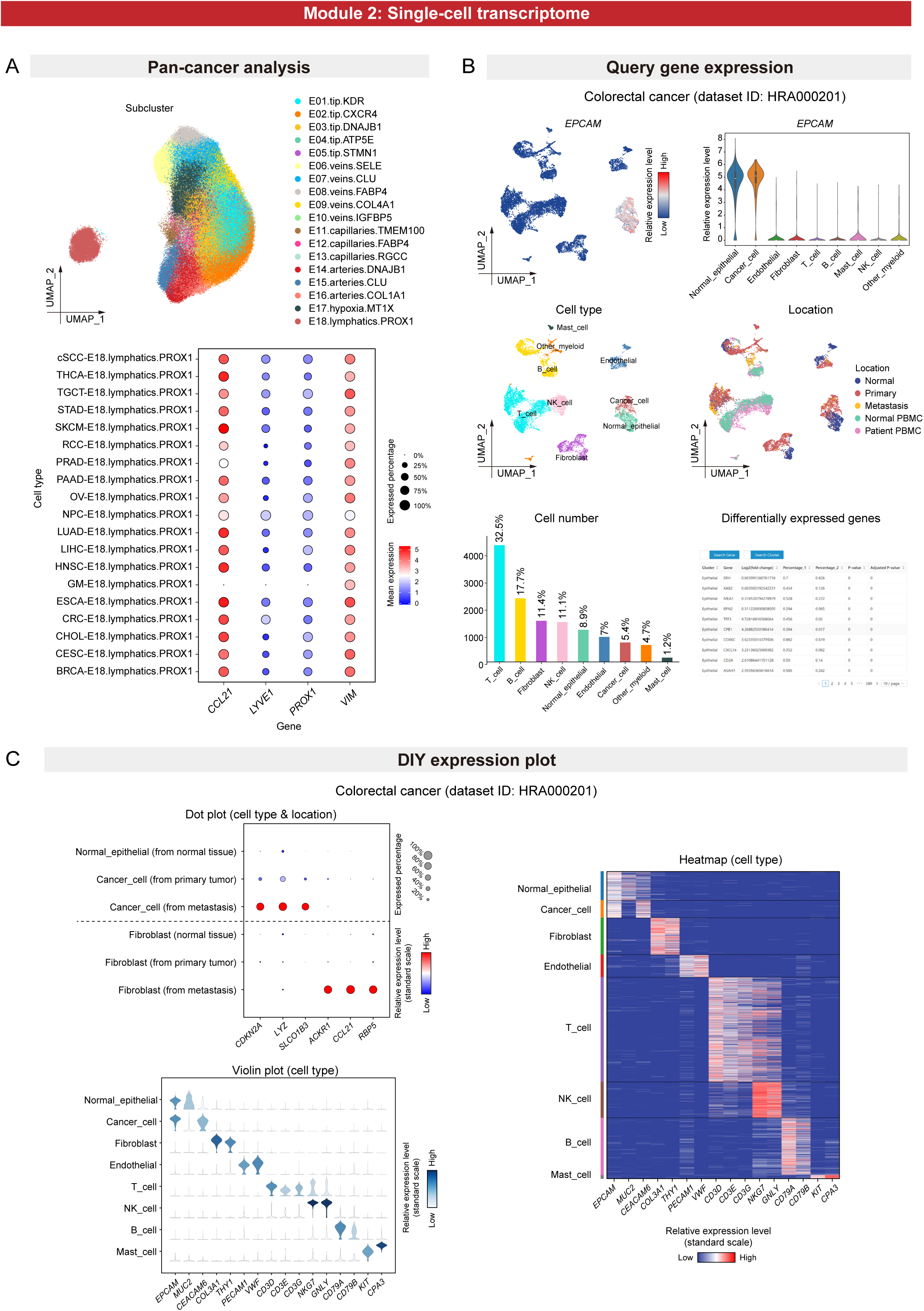
Examples for using the “Single-cell transcriptome” module. (A) The uniform manifold approximation and projection (UMAP) plot shows the subclusters of endothelial cells of human pan-cancer. The dot plot shows the expression levels and expressed percentages of the selected genes in the lymphatic endothelial cells (E18.lymphatics.PROX1). (B) With “Query gene expression” function, the relative expression levels of selected gene (*EPCAM*) across various cell types in colorectal cancer (dataset ID: HRA000201) are shown in UMAP plot and violin plot. Other related information, such as cell type identities, sampling locations, cell numbers and fractions of each cell type, and tables for differentially expressed genes, were also provided. (C) With the “DIY expression plot” function, the relative expression levels of multiple selected genes in multiple selected cell groups of colorectal cancer (dataset ID: HRA000201) are shown in three figure types, including dot plot, violin plot, and heatmap. For the dot plot, the expression levels of different cell types from different sampling locations (e.g. primary tumor v.s. metastasis) are plotted by selecting the “Cell type & location” options in the “Group by” select box. UMAP, uniform manifold approximation and projection. PBMC, peripheral blood mononuclear cell. Cancer types: cSCC, cutaneous squamous cell carcinoma; THCA, thyroid carcinoma; TGCT, testicular germ cell tumor; STAD, stomach adenocarcinoma; SKCM, skin cutaneous melanoma; RCC, renal cell carcinoma; PRAD, prostate cancer; PAAD, pancreatic adenocarcinoma; OV, ovarian cancer; NPC, nasopharyngeal cancer; LUAD, lung adenocarcinoma; LIHC, liver hepatocellular carcinoma; HNSC, head and neck squamous cell carcinoma; GM, glioma; ESCA, esophageal carcinoma; CRC, colorectal cancer; CHOL, cholangiocarcinoma; CESC, cervical squamous cell carcinoma and endocervical adenocarcinoma; BRCA, breast carcinoma.

#### Query gene expression

In addition to the above-mentioned “Pan-cancer analysis” function, users typically need to explore a certain cancer type in depth, which can be met by the “Query gene expression” function (Figure 3B). For example, users can easily explore the gene expression patterns of an interested gene (*EPCAM*) across different cell types of colorectal cancer, and get other related information at the same time, including the cell type identification, sample locations (e.g. normal tissue, primary tumor, and metastasis), cancer stages, fractions of each cell cluster, sex information, and the differentially expressed genes (DEGs) of each cell type (Figure 3B) (8). Moreover, a brief introduction and links to further information for each human gene are presented.

#### DIY expression plot

The “DIY expression plot” function enables users to query the expression levels of multiple selected genes in multiple selected cell types and generate customized figures for visualization (Figure 3C). Multiple figure types are provided, including dot plots, heatmaps, and violin plots, which can meet most of the needs of users. Moreover, for each cell type, users can easily compare the expression levels among different sampling locations (e.g. primary tumor v.s. metastasis), different cancer stages (e.g. stage I v.s. stage III), various cancer subtypes (e.g. poorly-differentiated adenocarcinoma v.s. well-differentiated adenocarcinoma), and between male and female by simply switching options in the “Group by” select box.

### Single-cell epigenome module

#### Focal DNA methylation plot

With this function, users can visualize the DNA methylation levels of a selected genomic region of interest (e.g. promoters) at single CpG resolution (Figure 4A). For example, the “Tanghulu” plot displayed the demethylated region in colorectal cancer cells compared with normal cells (Figure 4A) (7). Each row in the plot represents a cell group, with each circle denoting a CpG site.

**Figure 4.**
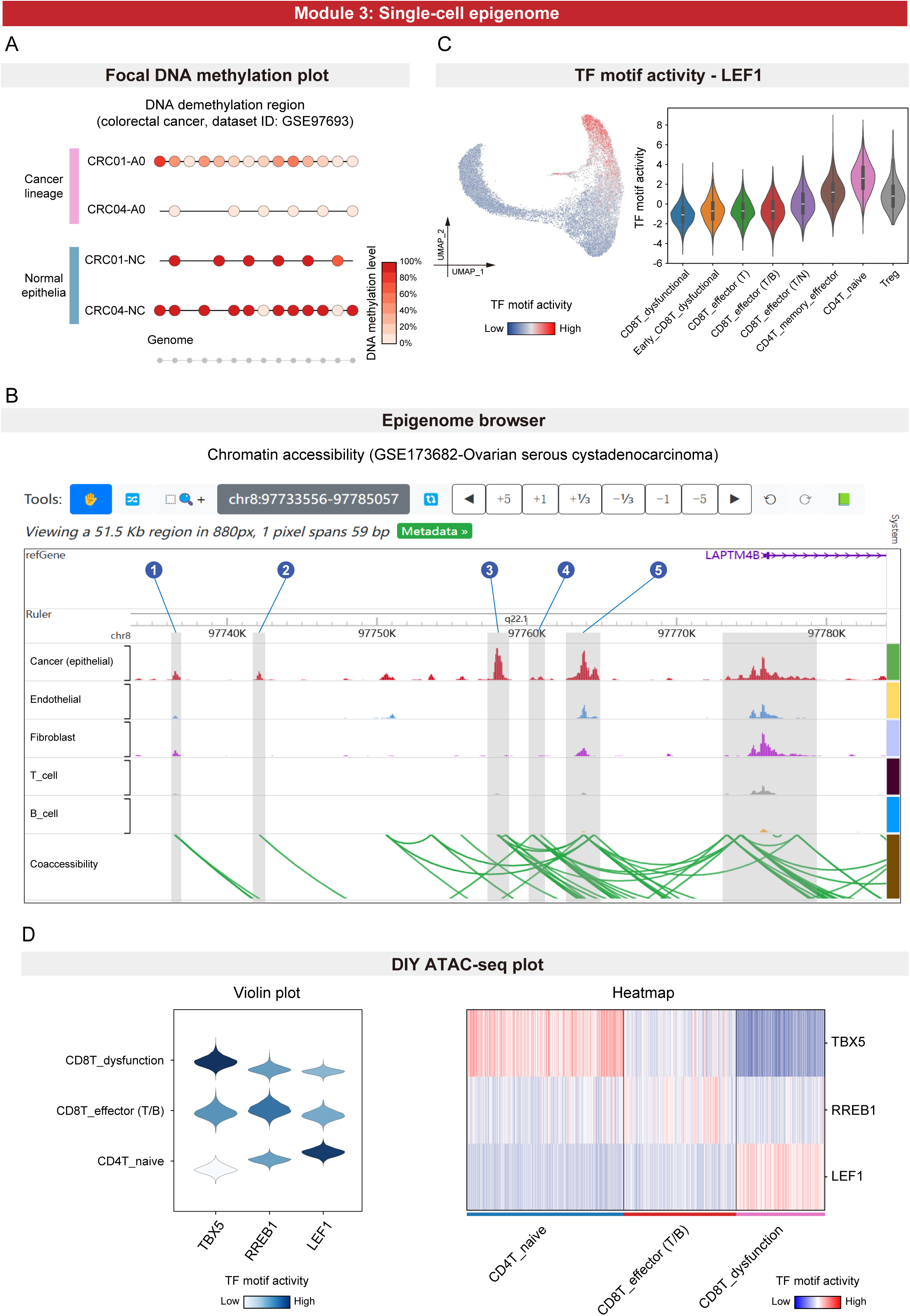
Examples for using “Single-cell epigenome” module. (A) The “Tanghulu” plot displays DNA demethylation regions in colorectal cancer cells (dataset ID: GSE97693), with each circle representing a CpG site. White dots represent unmethylated sites, while black dots represent methylated sites. Gray dots at the bottom indicate covered CpG sites within selected regions in selected cells. (B) Screenshot of the epigenome browser showing the chromatin accessibility levels around the transcription start site of the selected gene (*LAPTM4B*, Lysosomal Protein Transmembrane 4 Beta) of multiple cell types from ovarian cancer (dataset ID: GSE173682). The gray regions illustrate the potential enhancers in cancer cells. (C) Querying the motif activity of the transcription factor LEF1 across various cell types in kidney renal clear cell carcinoma (dataset ID: GSE181062) with the “TF motif activity” function. (D) Querying the motif activity of multiple transcription factor in multiple cell types in kidney renal clear cell carcinoma (dataset ID: GSE181062) with “DIY ATAC-seq plot” function.

#### Promoter DNA methylation plot

Users can query the DNA methylation levels of gene promoter regions at single-cell resolution, and compare them across normal tissues, primary tumors, and metastases with this module (Figure 2C).

#### Epigenome browser

scCancerExplorer incorporates the WashU Epigenome browser (32) to facilitate the exploration of chromatin accessibility and co-accessibility patterns (Figure 4B). Users can conveniently access specific genomic regions or genes of interest by inputting the official gene symbol or a genomic range in the search box at the top left of the browser interface. For example, we can see five potential enhancers for *LAPTM4B* (Lysosomal Protein Transmembrane 4 Beta) in ovarian cancer cells which showed higher accessibility levels compared to other cell types (Figure 4B) (12). The browser interface allows users to zoom in/out, navigate to designated regions, and customize display options such as display mode (bar plot or heatmap), RGB colors, axis scales, and text labels to suit individual preferences.

#### TF motif activity

With this function, users can search for an interested transcription factor and obtain the motif activity score (motif deviation z-score) across different cell types calculated using single-cell ATAC-seq data (Figure 4C).

#### DIY ATAC-seq plot

Similar to the “DIY expression plot” module, this module enables users to easily query the motif activities of multiple TFs across different cell types, and generate customized figures based on their requirements (Figure 4D).

### “Single-cell genome” module

This module allows users to investigate the single-cell copy number alterations (CNAs) of human cancers. For example, users can select a cancer type (e.g. stomach cancer) and select one or multiple patients (e.g. SC02 and SC07), and then generate the CNA heatmap (Figure 5A) (9). The cancer cells are sorted by genetic lineage if the lineage information is available.

**Figure 5.**
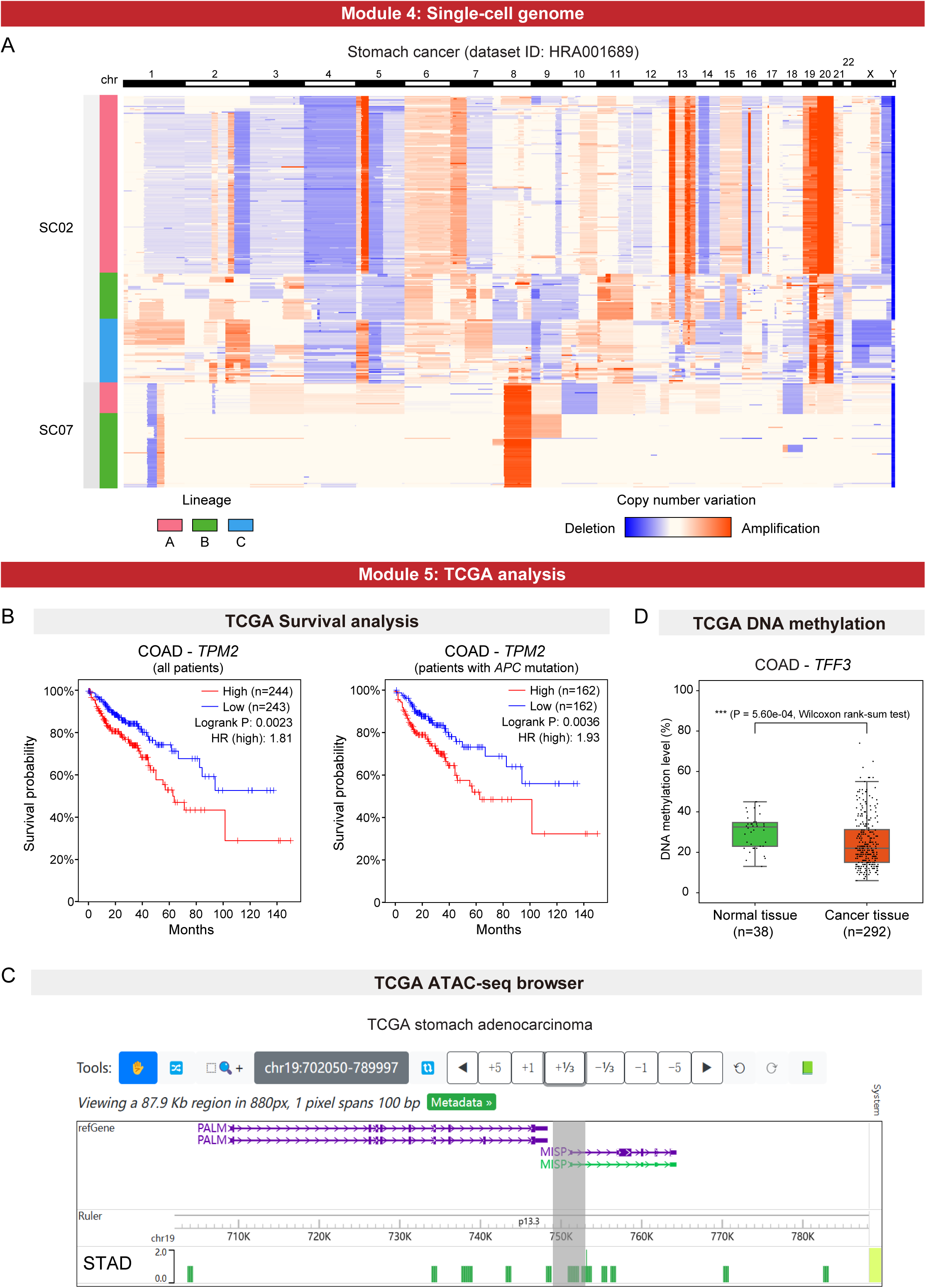
Examples for using “Single-cell genome” module and “TCGA analysis” module. (A) The heatmaps showing the copy number variations of stomach cancer patients (dataset ID: HRA001689). (B) All patients (left) or only the patients with *APC* mutation (right) in TCGA colon adenocarcinoma dataset were selected to perform survival analysis according to the expression levels of the selected gene (*TPM2*, Tropomyosin 2). (C) Browsing the chromatin accessibility pattern of the selected gene (*MISP*, Mitotic Spindle Positioning) in TCGA stomach adenocarcinoma ATAC-seq dataset with the “ATAC-seq browser” function. (D) The promoter DNA methylation levels of the selected gene (*TFF3*, Trefoil Factor 3) in normal tissues and cancer tissues in TCGA colon adenocarcinoma dataset. COAD, colon adenocarcinoma.

### “TCGA analysis” module

#### Survival analysis

This function provides survival analysis for 33 cancer types using TCGA data (Figure 5B). Notably, users can categorize patients based on multiple information, including gene expression levels (e.g. above median v.s. below median), gene mutation states (with mutation v.s. without mutation), pathological types (all types or certain subtype), and cancer stages (all stages or certain stage). For example, users can group patients by the expression levels of *TPM2* (Tropomyosin 2) in all colon adenocarcinoma patients or only in patients with *APC* (APC regulator of WNT signaling pathway) mutation, respectively (Figure 5B). Our previous study have identified *TPM2* (Tropomyosin 2) as fibroblast-specific biomarkers of poorer prognosis of colorectal cancer (8), which is consistent with the results.

#### ATAC-seq browser

With the “ATAC-seq browser” function, users can explore the open chromatin regions using TCGA ATAC-seq data. They can visualize the chromatin accessibility pattern of a gene of interest. For example, the open chromatin peaks around the promoter region of *MISP* (Mitotic Spindle Positioning), which are highly expressed in gastric cancer (9), can be browsed with this function (Figure 5C).

#### TCGA promoter DNA methylation

Users can query the DNA methylation levels of gene promoter regions for both normal tissues and tumor tissues using the TCGA DNA methylation data in this function (Figure 5D).

## DISCUSSION

The interplay between different molecular layers is critical for cancer progression and metastasis. Integrated single-cell multi-omics analyses are valuable for unraveling the complexities of cancer biology and facilitating the development of personalized therapeutic strategies. To the best of our knowledge, scCancerExplorer is the most comprehensive database for exploring single-cell multi-omics data of human pan-cancer currently.

First, by preprocessing a large number of single-cell multi-omics data and providing rich interactive tools for exploring them, we provide an important bridge between the valuable but hard-to-process resources and cancer researchers. The 5 major modules and 12 useful functions meet most of the requirements of users for exploring both single-cell genome, epigenome, and transcriptome data, and TCGA data.

Second, our featured module “Integrated multi-omics analysis” integrates multi-omics data of each gene on one web page, including TCGA bulk RNA-seq data, single-cell RNA-seq data, TCGA DNA methylation data, single-cell DNA methylation data, single-cell chromatin accessibility data, and TCGA survival analysis results. This module provides comprehensive and convenient knowledge for users to learn an interested gene from multiple perspectives, and helps users gain clues for further investigations.

Third, several customizable functions and publishing-ready figures were provided to meet the personalized requirements of users. For example, with the “DIY expression plot” function, users can select multiple genes and cell types and designate their order shown in the generated figures, download the editable PDF files, and use the publishing-level figures in their study. Additionally, by simply switching options in the “Group by” select box, users can easily compare the expression levels between primary tumors and metastases, pre-treatment and post-treatment tumors, and various cancer subtypes.

Fourth, not only single-cell multi-omics data, but also TCGA multi-omics data were collected and analyzed, including bulk RNA-seq data, DNA methylation 450K data, and bulk ATAC-seq data. Moreover, we provide customizable TCGA survival analysis, which allows users to group patients not only by the expression levels of the selected gene but also by gene mutation states (e.g., *TP53* mutations), pathological subtypes and cancer stages.

## DATA AVAILABILITY

The database is available at https://bianlab.cn/scCancerExplorer.

## FUNDING

This work was supported by the National Natural Science Foundation of China (grant numbers 82173059, 82172956), the Young Elite Scientists Sponsorship Program by China Association for Science and Technology (CAST) (grant number 2020QNRC001), and research start-up funding from Nanjing Medical University (grant number KY116RC20200007).

## CONFLICT OF INTEREST

The authors declare that they have no competing interests.

## ACKNOWLEDGEMENTS

We gratefully acknowledge Prof. Jingchu Luo (Peking University) for giving advices for constructing databases. We gratefully thank Prof. William Greenleaf (Stanford University) and his group for providing help when processing single-cell ATAC-seq data. Part of the analysis was conducted using the high-performance computing platform at Nanjing Medical University. We gratefully acknowledge the assistance of Tao Wang from the Department of Information Technology Construction and Management at Nanjing Medical University.

## Notes

https://bianlab.cn/scCancerExplorer

### Competing Interest Statement

The authors have declared no competing interest.

### Summary of Updates

Figures 1-5 were revised. Manuscript was revised.

